# Chess expertise improves rule-guided flexibility and visual working memory precision

**DOI:** 10.64898/2026.06.28.733231

**Authors:** Marjan Makhsous, Mohammad Jowkar, Ehsan Rezayat

## Abstract

Studying chess experts helps researchers understand how intensive practice shapes thinking skills. Cognitive flexibility is the ability to adjust thoughts when rules or tasks change. Working memory is the ability to hold and use information over short periods. This study compared cognitive flexibility and working memory precision between adolescent chess players and non-players. Twenty-four professional chess players and twenty-five controls completed two novel behavioral tasks. Chess players showed better accuracy in both tasks than controls. They adapted more efficiently when rules changed during a continuous learning task. They also remembered facial expressions more precisely in a working memory task. Learning rates in the flexibility task did not differ between groups. These results indicate that chess expertise may improve rule-guided flexibility and visual working memory precision in adolescents.

## Introduction

Chess is more than a game. It requires holding complex board states in mind, revising strategies as the opponent moves, and switching between rules moment by moment. Neuroimaging studies show that chess expertise engages frontoparietal and visuospatial networks efficiently. Experts also show reduced cognitive cost during problem solving (Williams et al., 2025). Other domains of expertise, such as music and medical decision-making, suggest that long-term structured practice refines neural systems for high-level cognition (Criscuolo et al., 2022). Together, these results indicate that chess offers a natural model to study how experience shapes executive functions.

Chess engages several executive functions. Two of the most important are cognitive flexibility and working memory. Cognitive flexibility is the ability to adjust thoughts and actions when task demands change. It supports adaptive behavior. Yet its definition is not always clear. Different tasks measure different aspects – task switching, reversal learning, attentional shifting. Some focus on rule changes, others on probabilistic feedback. This study focuses on rule-guided flexibility: how well people adapt when explicit rules reverse. This form of flexibility relies on frontoparietal and midcingulo-insular networks (Kupis & Uddin, 2023) and rapid changes in prefrontal-parietal connectivity (Cole, 2024). Working memory is the other core component. It holds task-relevant information over short periods. Contemporary models describe working memory as a noisy system. Precision – not just correct/incorrect – reflects the quality of stored information (Brady et al., 2024). Precision varies with neural noise and sensory recruitment. Studies using continuous delayed-reproduction tasks show that precision is sensitive to load and experience (Esfahan et al., 2025). Binary tasks hide this nuance.

Although flexibility and working memory are distinct, they interact during rule maintenance, updating, and rapid adjustment. Both rely on prefrontal-parietal circuits. Both benefit from experience-dependent refinement. The present study examines rule-guided flexibility and visual working memory precision in adolescent chess players and matched controls. We used two novel continuous-error tasks – one for probabilistic reversal learning (CPRLT) and one for face-based working memory (FWMT). We asked whether chess expertise improves these specific components beyond general reinforcement learning.

## Method

### Participants

In this study 49 adolescents aged 13-19 years were recruited: 24 chess players and 25 non-chess players (control group). All chess players possessed significant professional experience, having engaged in the game for more than 4 years, and had participated in both national and international tournaments. All participants reported normal or corrected-to-normal vision and no history of neurological or psychiatric disorders. Written informed consent was obtained from legal guardians, and the study protocol was approved by the University of Tehran Science Committee (IR.UT.PSYEDU.REC.1404.064). Participants completed two tasks as described below during a single session, with the order of administration counterbalanced across individuals. All testing was conducted in a quiet, dimly lit room under standardized conditions, with participants seated approximately 60 cm from the monitor.

### Tasks

#### Face Working Memory Test (FWMT)

This task assessed the precision of visual working memory for facial stimuli and followed procedures similar to those used in recent works on face-based memory precision (Asgari, A., Vahabie, A. H., & Rezayat, 2025; Haghian et al., 2025). The task consisted of three blocks with 54 trials each (see Figure 1B). Each trial began with a central fixation cross (1 s), followed by the sequential presentation of three sample faces (1 s each). After a retention interval of either 1.5 or 3 s, a numerical cue indicated which of the three faces should be recalled.

**Figure 1.**
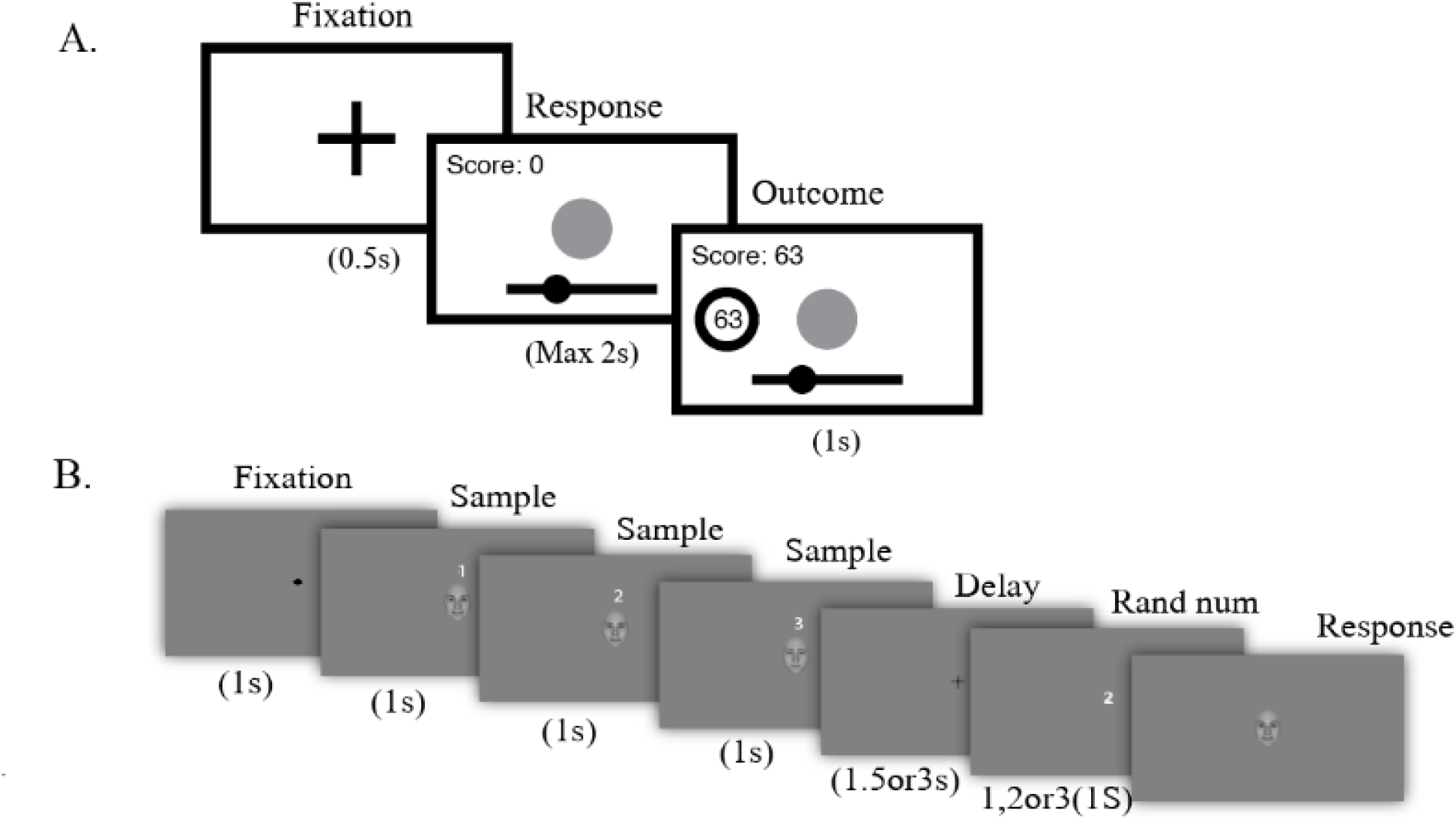
A) Continuous-score Probabilistic reversal Learning Test (CPRLT). Subjects change the size of the circle in the center of the screen by moving the mouse across the horizontal line at the bottom of the screen, and select a size by clicking the mouse. They then see a score between -100 to 100. They should try to find sizes that give them higher scores. B) Face Working Memory Task – trial structure. Subjects see three faces with different emotional expressions (morphed from sad to happy, The faces shown are computer-generated and do not represent any real individual.), each for 1 second. After a variable delay, they are required to reconstruct one of the previous faces, specified with its number.

Participants adjusted a test face along a morph continuum to match the cued sample. Stimuli were generated using FaceGen Modeller and luminance-matched with the SHINE toolbox in MATLAB. The task was programmed in PsychToolbox. Performance indices included mean absolute error (MAE), defined as the absolute difference between the reproduced and target morph index, and reaction time (RT). Since MAE data was bounded between -18 to +18, we fitted a von-Mises distribution to the MAE data (Bays, Catalao, & Husain, 2009), which results in a Kappa parameter that shows concentration of the distribution; higher Kappa shows better performance(Bays et al., 2009). After simulating random responding in this task for 1000 times, we found that Kappa value for random response was approximately 0.78. Any participant with Kappa lower than 0.78 was considered outlier.

#### Continuous-score Probabilistic Reversal Learning Test (CPRLT)

Cognitive flexibility was measured with a novel test acquired from the work of Jowkar (2025). This test consists of 10 training trials and 150 experimental trials. On each trial, participants adjusted the diameter of a green circle along a horizontal axis to approximate a hidden target size within 2000 ms (see Figure 1.A). Feedback was provided as a score ranging from –100 to +100. In some trials, small sizes were better, and in some trials, large sizes were better. This rule was reversed every 30 trials (at trials 31, 61, 91, and 121). Responses were scaled in the range of 0–1. The primary measure was mean absolute error (MAE) relative to the hidden optimal size. Participants with missed trials more than 10%, and participants with MAE higher than “2.5 × 1/12” (approximately 0.2) were classified as outliers. We also calculated an adjusted MAE score which focused on trials 31–150 to capture cognitive flexibility more accurately. Reinforcement learning processes were modeled using a simplified Rescorla–Wagner framework (Rescorla & Wagner, 1972). The model assumes that the subject has an expectation about the best scoring value (V) in each trial. The difference between the hidden value (λ) and V, is the prediction error(δ):

δt=λt−Vt

λt: Observed outcome at trial t; The hidden size in trial t

Vt: Current expected value; The subject’s belief about the best scoring size before trial t

The subject updates his/her expectation (V) according to the prediction error(δ), based on the following equation:

Vt+1=Vt+α⋅δt

α is the learning rate, which is the only free parameter of this model, and is between 0 and 1. Learning rate shows how quickly the subject updates his or her expectation(V): higher α shows quicker updates and vice versa. This parameter is not directly related to cognitive flexibility, but we believe that very low learning rate indicates lower levels of cognitive flexibility. We used maximum likelihood estimation to estimate α for every subject.

## Results

We recruited 24 chess players (13 male, 11 female) and 25 non-chess controls (12 male, 13 female). A chi-square test showed no significant difference in gender distribution between groups, χ^2^(1) = 0.19, p = 0.67. Age was comparable between groups (chess players: M = 15.96 years, SD = 1.76; controls: M = 15.96, SD = 1.65; t (47) = 0.12, p = 0.903). Normality was assessed using the Kolmogorov-Smirnov test; age deviated from normality (p < 0.001), but the remaining variables were approximately normally distributed (all p > 0.05; see Table 1).

**Table 1.**
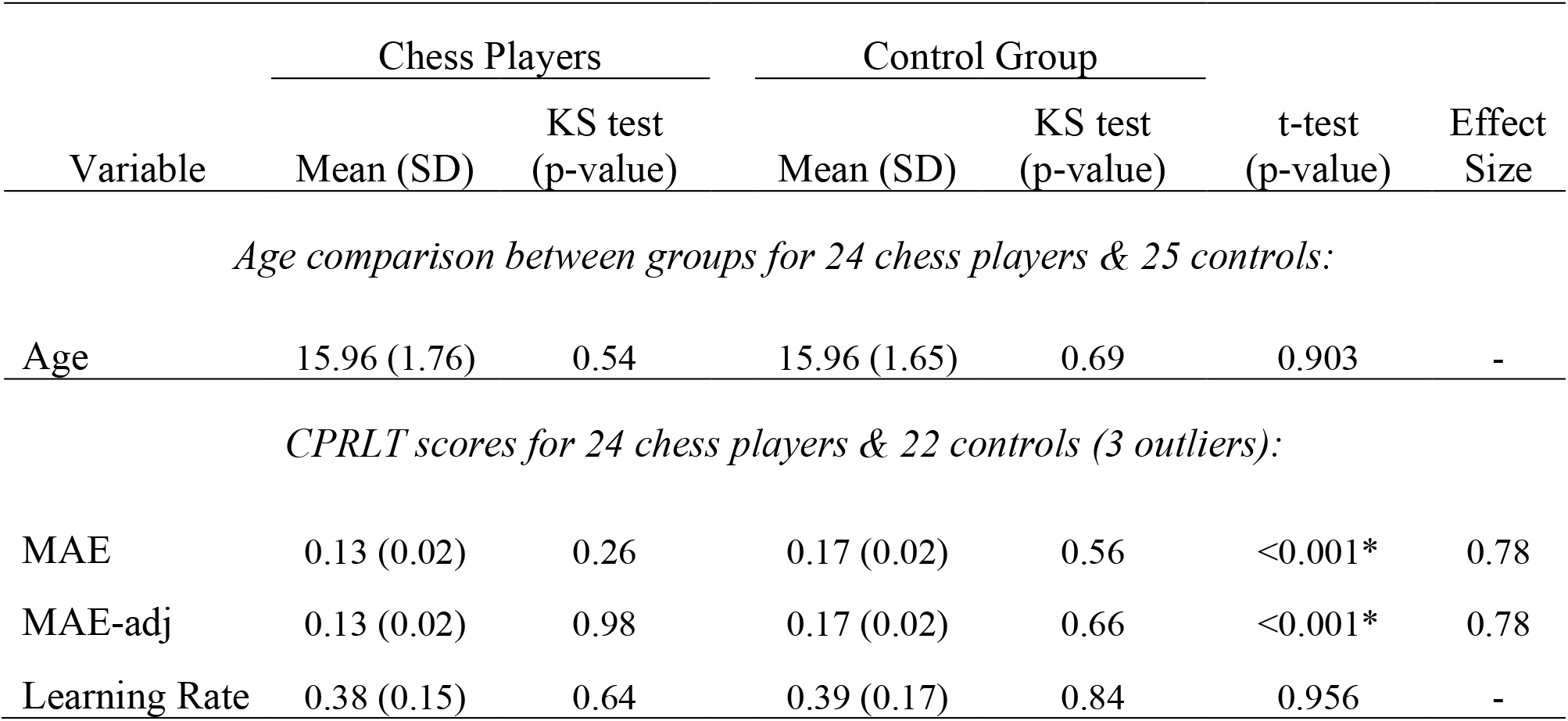
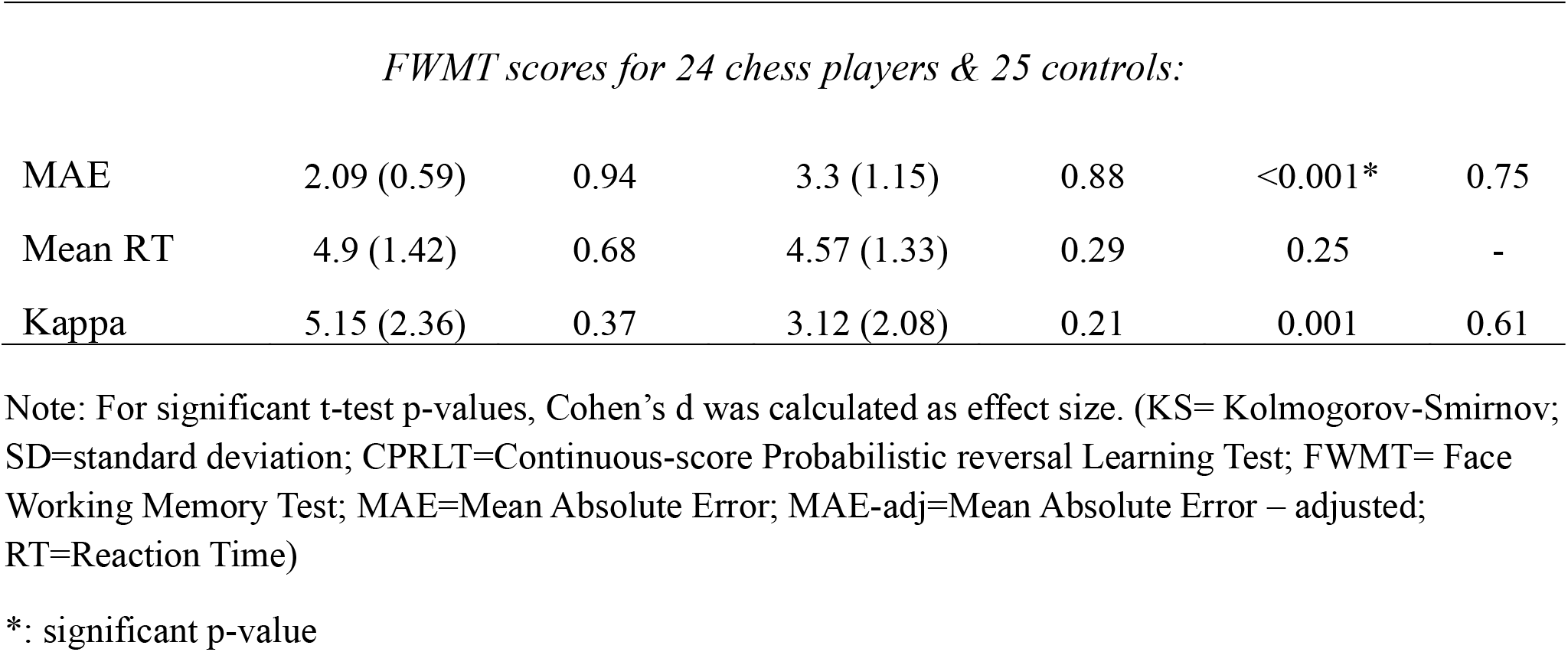
Descriptive statistics, Kolmogorov-Smirnov test result, and comparison of variables.

| Variable | Chess Players |  | Control Group |  | t-test<br>(p-value) | Effect<br>Size |
| --- | --- | --- | --- | --- | --- | --- |
|  | Mean (SD) | KS test<br>(p-value) | Mean (SD) | KS test<br>(p-value) |  |  |
| Age comparison between groups for 24 chess players & 25 controls: |  |  |  |  |  |  |
| Age | 15.96 (1.76) | 0.54 | 15.96 (1.65) | 0.69 | 0.903 | - |
| CPRLT scores for 24 chess players & 22 controls (3 outliers): |  |  |  |  |  |  |
| MAE | 0.13 (0.02) | 0.26 | 0.17 (0.02) | 0.56 | <0.001* | 0.78 |
| MAE-adj | 0.13 (0.02) | 0.98 | 0.17 (0.02) | 0.66 | <0.001* | 0.78 |
| Learning Rate | 0.38 (0.15) | 0.64 | 0.39 (0.17) | 0.84 | 0.956 | - |
| *FWMT scores for 24 chess players & 25 controls:* |  |  |  |  |  |  |
| MAE | 2.09 (0.59) | 0.94 | 3.3 (1.15) | 0.88 | <0.001* | 0.75 |
| Mean RT | 4.9 (1.42) | 0.68 | 4.57 (1.33) | 0.29 | 0.25 | - |
| Kappa | 5.15 (2.36) | 0.37 | 3.12 (2.08) | 0.21 | 0.001 | 0.61 |

Note: For significant t-test p-values, Cohen's d was calculated as effect size. (KS= Kolmogorov-Smirnov; SD=standard deviation; CPRLT=Continuous-score Probabilistic reversal Learning Test; FWMT= Face Working Memory Test; MAE=Mean Absolute Error; MAE-adj=Mean Absolute Error – adjusted; RT=Reaction Time)
\*: significant p-value

In the FWMT, no outliers were detected. Chess players showed higher working memory precision than controls. Their MAE was lower (M = 2.09, SD = 0.59) compared to controls (M = 3.30, SD = 1.15), t (47) = 4.57, p < 0.001, d = 0.75, Figure 2 and Figure 4). The concentration parameter (kappa) was also higher for chess players (M = 5.15, SD = 2.36) than controls (M = 3.12, SD = 2.08), t (47) = 3.27, p = 0.001, d = 0.61. Mean reaction time did not differ significantly between groups (chess players: M = 4.90 s, SD = 1.42; controls: M = 4.57 s, SD = 1.33; t (47) = 1.17, p = 0.25).

**Figure 2.**
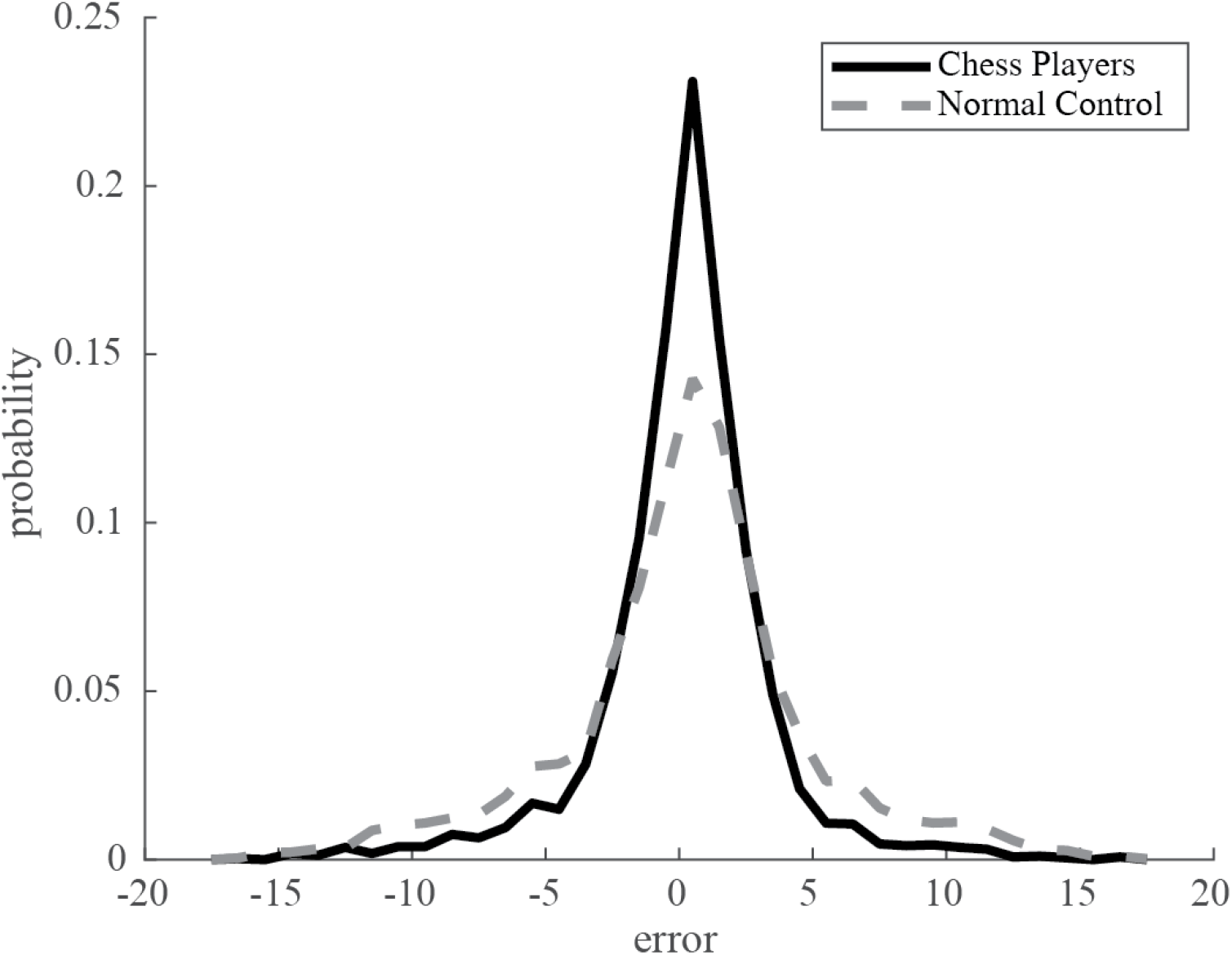
Overall plot for error distribution in Face Working Memory Task (FMWT). For each distribution, all trials of all participants of the group were used together

In the CPRLT, three control participants were excluded as outliers because their mean absolute error (MAE) exceeded the acceptable threshold of 0.2. The final analysis included 24 chess players and 22 controls. Chess players performed significantly better than controls on both rule-based flexibility measures. Their MAE was lower (M = 0.13, SD = 0.02) compared to controls (M = 0.17, SD = 0.02), t (44) = 6.72, p < 0.001, Cohen’s d = 0.78. Similarly, the adjusted MAE (MAE_adj) was lower for chess players (M = 0.13, SD = 0.02) than controls (M = 0.17, SD = 0.02), t (44) = 6.70, p < 0.001, d = 0.78, Figure 3, and figure 4). However, the learning rate (alpha), which indexes reward-based free learning, did not differ between groups (chess players: M = 0.38, SD = 0.15; controls: M = 0.39, SD = 0.17; t (44) = 0.06, p = 0.956).

**Figure 3.**
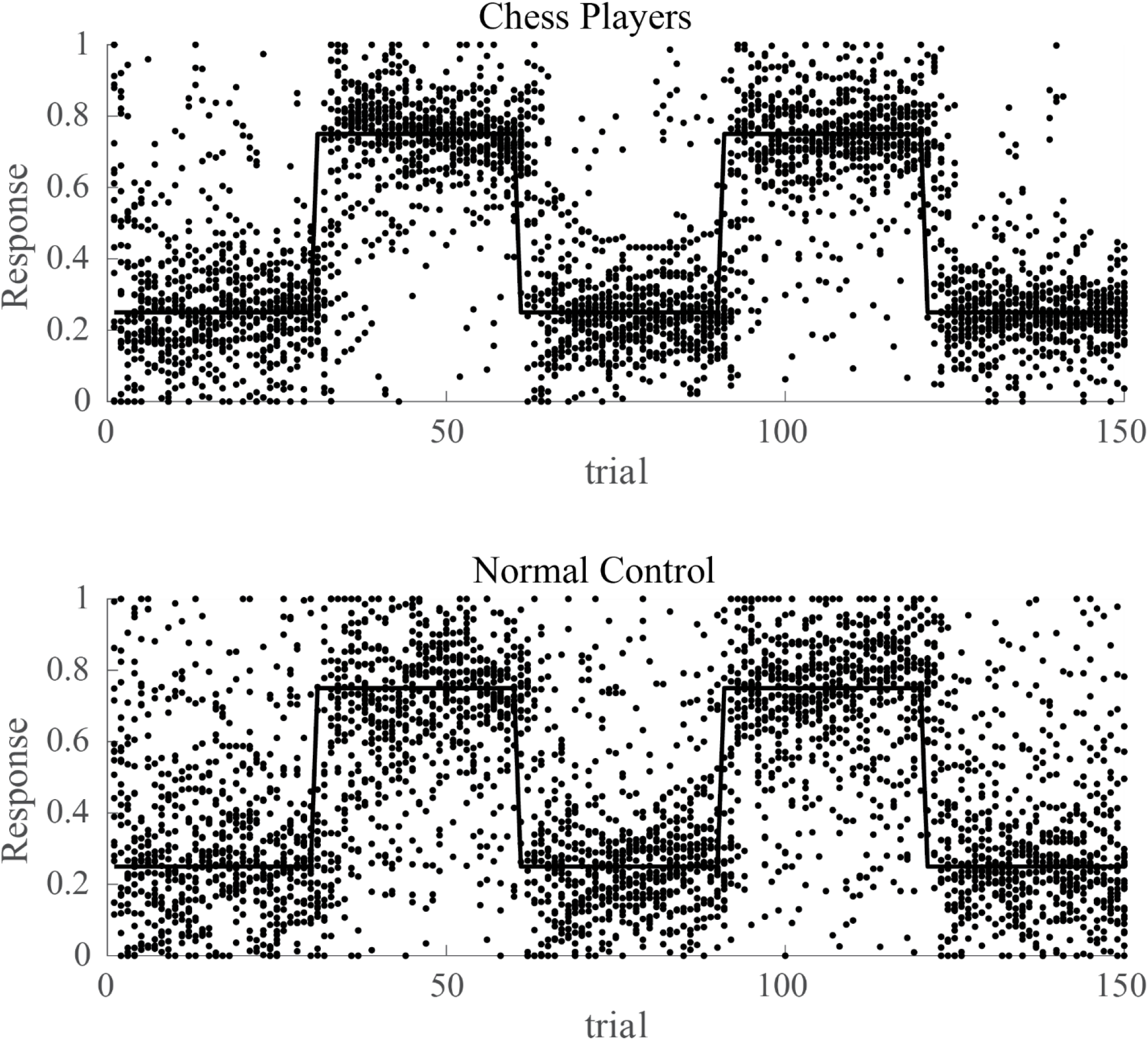
Overall plot for Continuous-score Probabilistic Reversal Learning Test (CPRLT). Selected responses for all chess players (panel A) and normal controls (panel B) are plotted across trials. As can be seen, both groups selected sizes around the optimal size (indicated by the line plot), but chess players selected more accurately and their responses has lower variance around the optimal size, compared to normal controls. This indicates similarities in general reinforcement learning, while differences rule-guided learning among groups.

**Figure 4.**
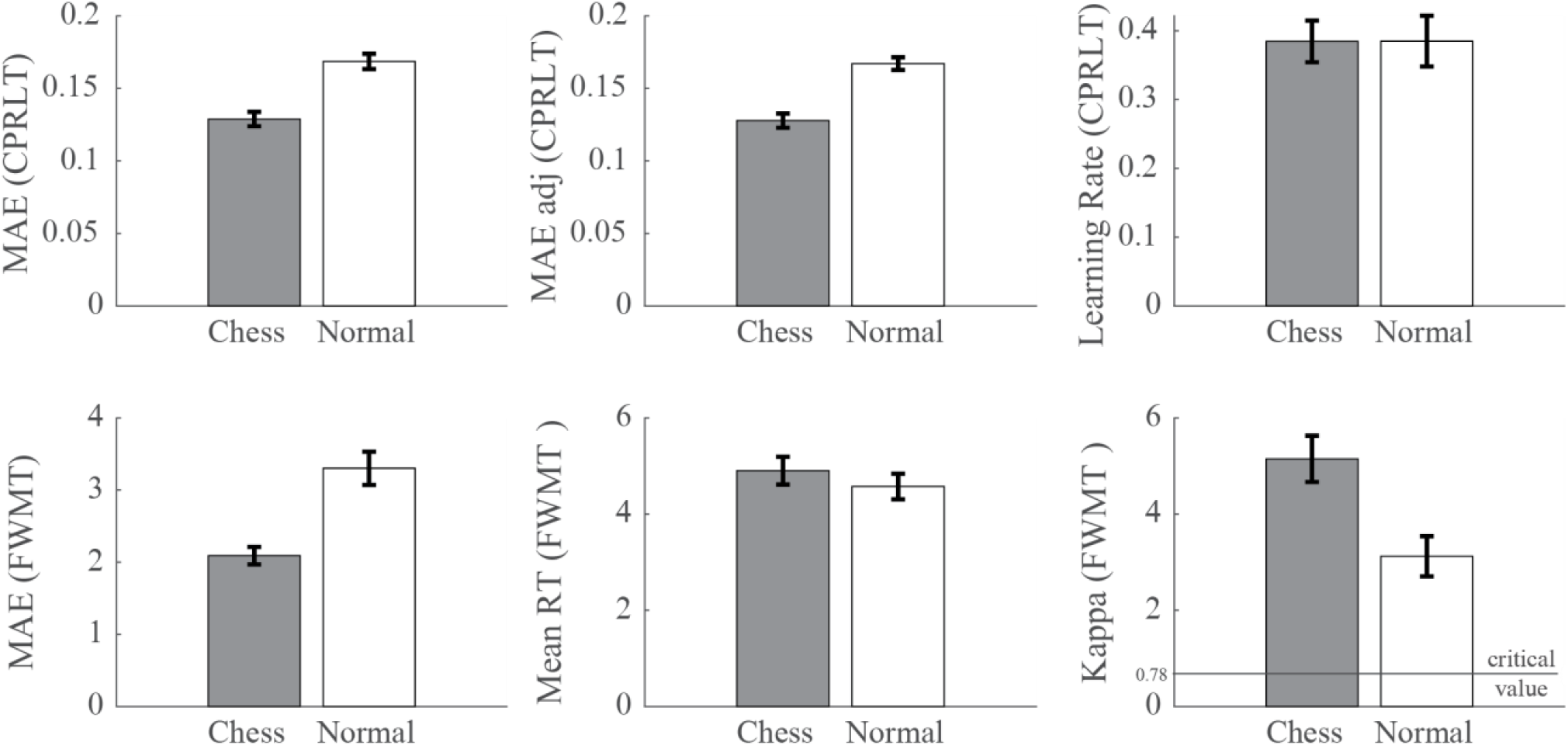
Comparison of scores among chess players and normal subjects. Y-axis shows the mean, and error bars show the standard error of mean. Except for the “Learning Rate (CPRLT)” and “Mean RT (FMWT), all differences are statistically significant.

These results indicate that chess expertise is associated with better rule-guided flexibility and more precise face-based working memory, but not with faster learning from probabilistic feedback or faster reaction times.

## Discussion

The present study introduces two novel task designs that extend current approaches to executive function research. First, the cognitive flexibility task measured performance on a continuous error scale rather than binary correct–incorrect outcomes. This design provides a more sensitive index of adaptive learning, capturing subtle variations in rule updating that are often missed in categorical measures (Cole, 2024). Second, the working memory task employed facial stimuli, building on recent evidence that memory precision for socially relevant stimuli can reveal distinct patterns compared to non-social features (Esfahan, S. M., Sepahi, N., & Rezayat, 2025). By combining a continuous measure of flexibility with a face-based working memory paradigm, the study offers a methodological advance that allows for finer analysis of executive processes in adolescents.

In the cognitive flexibility task, chess players showed superior performance in rule-based learning. They adapted more efficiently when explicit rules guided behavior, consistent with the idea that structured practice enhances the ability to detect and apply regularities. By contrast, no group differences emerged in free learning, where adaptation relied only on probabilistic reward feedback. This pattern suggests that chess expertise may selectively benefit rule-guided flexibility rather than general reinforcement learning (figure 3). Prior work has emphasized that rule learning engages prefrontal circuits involved in maintaining and updating structured representations (Kupis, Lauren B.; Uddin, 2023), whereas free learning depends more on trial-by-trial reward history. The present findings extend this distinction by showing that adolescents with chess expertise gain an advantage specifically in the rule-based domain.

In the face working memory task, chess players demonstrated higher accuracy and lower error compared to controls. This advantage indicates that expertise may enhance the fidelity of visual memory for socially relevant stimuli, consistent with prior findings that precision is a sensitive marker of working memory quality (Asgari, A., Vahabie, A. H., & Rezayat, 2025).

Research on chess has long examined its links to executive processes, including planning, attention, and working memory. Studies of expert players show enhanced recruitment of frontoparietal and visuospatial networks during problem solving, with reduced cognitive cost compared to novices (Williams, L. H.; Störmer, V. S.; Brady, 2025). Other domains of expertise, such as music and medical decision-making, similarly demonstrate that structured practice can refine neural systems supporting high-level cognition (Criscuolo A, Schwartze M, 2022)(Cera N, Pinto J, Dong M, Durning S, 2025). Within this literature, chess has been proposed as a natural model for studying how sustained practice shapes executive functions. However, most prior work has focused on broad cognitive outcomes, such as general intelligence or attentional control, rather than specific computational components. The present study extends this line of research by showing that chess expertise relates to rule-guided flexibility and to the precision of working memory for faces. These findings suggest that chess may provide a useful framework for probing the mechanisms of adaptive learning and memory fidelity in adolescence.

This study has several limitations. The sample was restricted to adolescents, and only two task paradigms were used. Future work should examine whether similar patterns emerge across different age groups and with a broader range of executive tasks. Despite these constraints, the findings provide converging evidence that chess expertise relates to rule-guided flexibility and to the precision of working memory for faces. Together, the results suggest that structured practice in chess may serve as a useful framework for probing how experience shapes core executive functions during development.

## Conclusion

The study examined cognitive flexibility and working memory precision in adolescent chess players using two novel task designs. Results showed that chess expertise was linked to better performance in rule-guided flexibility and to higher accuracy in face-based working memory. These findings suggest that structured practice in chess may selectively strengthen executive processes that rely on rule detection and memory fidelity. While the sample was limited to adolescents and only two paradigms were tested, the results provide converging evidence that chess can serve as a natural model for investigating how sustained practice shapes core executive functions. Future research should extend this approach to broader populations and task domains to clarify the mechanisms through which expertise influences adaptive cognition.

